# An Investigation of Parameter-Dependent Cell-Type Specific Effects of Transcranial Focused Ultrasound Stimulation Using an Awake Head-Fixed Rodent Model

**DOI:** 10.1101/2024.06.24.600515

**Authors:** Sandhya Ramachandran, Huan Gao, Eric Yttri, Kai Yu, Bin He

## Abstract

Transcranial focused ultrasound (tFUS) is a promising neuromodulation technique able to target shallow and deep brain structures with high precision. Previous studies have demonstrated that tFUS stimulation responses are both cell-type specific and controllable through altering stimulation parameters. Specifically, tFUS can elicit time-locked neural activity in regular spiking units (RSUs) that is sensitive to increases in pulse repetition frequency (PRF), while time-locked responses are not seen in fast spiking units (FSUs). These findings suggest a unique capability of tFUS to alter circuit network dynamics with cell-type specificity; however, these results could be biased by the use of anesthesia, which significantly modulates neural activities. In this study, we develop an awake head-fixed rat model specifically designed for tFUS study, and address a key question if tFUS still has cell-type specificity under awake conditions. Using this novel animal model, we examined a series of PRFs and burst duty cycles (DCs) to determine their effects on neuronal subpopulations without anesthesia. We conclude that cell-type specific time-locked and delayed responses to tFUS as well as PRF and DC sensitivity are present in the awake animal model and that despite some differences in response, isoflurane anesthesia is not a major confound in studying the cell-type specificity of ultrasound neuromodulation. We further determine that, in an awake, head-fixed setting, the preferred PRF and DC for inducing time-locked excitation with our pulsed tFUS paradigm are 1500 Hz and 60%, respectively.

## Introduction

Neuromodulation has received considerable attention as a non-pharmacologic treatment of neurological disorders such as Alzheimer’s disease, Parkinson’s disease, epilepsy, depression, anxiety, chronic pain, and more. To treat these neurological disorders, precise stimulation of specific regions of brain and neural circuits is required. Neuromodulation techniques, in particular those based on electromagnetic stimulation principles have been widely studied (Ashkan et al., 2017; Berlim et al., 2013; Bikson & Rahman, 2013; Chail et al., 2018; Filmer et al., 2014; Gardner, 2013; Giordano et al., 2017; Horvath et al., 2011; M. D. Johnson et al., 2013; N. N. Johnson et al., 2018; Klomjai et al., 2015; Koponen & Peterchev, 2020; Lozano et al., 2019; ME et al., 2014; Nitsche et al., 2008, 2009; Nitsche & Paulus, 2000; Roy et al., 2014; Thair et al., 2017), but have limitations due to invasiveness, side effects, or ability to precisely stimulate the brain region or penetrate into the deep brain structures.

Transcranial focused ultrasound (tFUS) is a noninvasive technology for modulating brain function with focused ultrasound pressure waves (Blackmore et al., 2019; Fini & Tyler, 2017; Kamimura et al., 2020; Naor et al., 2016; Tufail et al., 2010, 2011). It has a spatial resolution of a few millimeters in humans and as high as 100 µm in mice (Cheng et al., 2022; Kamimura et al., 2020), as well as the ability to target at deep brain structures (Dallapiazza intrinsic et al., 2018; Folloni et al., 2019; Legon et al., 2018). Additionally, recent studies have indicated that it has cell-type specific effects controllable through parameter selection (Plaksin et al., 2016; Yu et al., 2021), making it a promising neuromodulation modality.

Although these findings are promising, previous studies relied on using anesthetized rodent models, confounding these results. The anesthetized brain functions differently than any awake or asleep brain state, with observed modulation of hemodynamics, metabolic state, receptive field size, fMRI resting state network, and overall reduction of neural activity and responsivity (Gao et al., 2017). Isoflurane anesthesia in particular is known to act on potassium channels and calcium channels, as well as on the GABA receptor complex present in inhibitory neurons (Buljubasic et al., 1992; Kokita et al., 1999; Moody et al., 1993; Thrane et al., 2012). Several theories (Moody et al., 1993) of tFUS’s mechanism involve modulation of ion channel properties, which these anesthesia effects would likely interact with (Chail et al., 2018; Krasovitski et al., 2011; Yoo et al., 2022). Physiological changes from anesthesia may also impact the neural effects of tFUS, including changes in the volume fraction of extracellular space, brain temperature, intracranial pressure, and isoflurane’s opening of the blood-brain barrier (Artru, 1984; Gao et al., 2017; Tétrault et al., 2008). Anesthesia effects also result in more widely varying responsivity to stimulus over time under its effects, potentially resulting in some of the high variance of responses to tFUS that are observed (Castro-Alamancos, 2009; Curto et al., 2009; Gao et al., 2017; Liang et al., 2015; Sellers et al., 2013, 2015; Steriade et al., 2001).

Of particular interest is whether anesthesia is responsible for our previous finding of cell type specificity, as well as parameter sensitivity. (H. Kim et al., 2021; King et al., 2013a; Lee et al., 2015)(Dallapiazza et al., 2018; Niu et al., 2022; Ramachandran et al., 2022; Sanguinetti et al., 2020; Verhagen et al., 2019)Recent work by our group as well as others have demonstrated that parameters such as the pulse repetition frequency (PRF) and the duty cycle (DC) are particularly significant when it comes to tuning the amount of tFUS effect (H. Kim et al., 2014; King et al., 2013b; Manuel et al., 2020; Yu et al., 2021). The effects of these parameters appear to vary between excitatory and inhibitory neurons in the cortex (Murphy et al., 2022a; Yu et al., 2021), studied through sorting recorded neuronal spikes into fast spiking units (FSU) and regular spiking units (RSU), which are putative inhibitory interneurons and excitatory pyramidal neurons, using their action potential waveform shapes (Cauli et al., 1997; Geiger et al., 1995; Martina et al., 1998; Martina & Jonas, 1997; Morin et al., 1996; Murray & Keller, 2011; Savić et al., 2001; Taverna et al., 2005). Specifically, our group observed that in the sensory cortex, only RSU have a time-locked response to tFUS that is sensitive to the change of PRF. This finding implies that tFUS may be able to modulate dynamics between excitatory and inhibitory cells through careful choice in stimulation parameters, opening possibilities for a variety of therapeutic treatments. However, these cell types are also differently affected by anesthesia, since as mentioned before, anesthesia selectively acts on receptors in inhibitory cells to reduce consciousness (Moody et al., 1993). Anesthesia has also been shown to reduce thalamo-cortical connectivity in rats, which may specifically confound our findings in S1 (Gao et al., 2017; Liang et al., 2012, 2013; Liu et al., 2011). Due to these confounds, it is critical to test these findings in an awake model.

Ultrasound has been tested in awake rodents before, but most commonly it is delivered under anesthesia and effects tested after awakening, which only allows for testing of long-term effects (M. G. Kim et al., 2022). Wearable transducers for rodents have been developed, but generally only allow for EEG recordings to be taken due to the higher noise levels from movement, and these transducers are not capable of as high precision or as wide a range of parameters (Di Ianni et al., 2023; Hou et al., 2024; Jo et al., 2022; Piech et al., 2017). To target the brain with tFUS precisely in an awake animal while taking high frequency recordings, head-fixation is required to keep the animal still. However, awake head-fixation of rats for tFUS poses several unique challenges. In general, head fixation involves surgically attaching a headpiece to an animal’s skull using adhesives and screws, which after recovery can be locked into a set- up that keeps the animal still. This is typically done with mice because they are smaller and much less strong, requiring less training (Gao et al., 2017; Schwarz et al., 2010). For tFUS however, a rat model with larger skull size is preferred, providing enough space to insert an electrode for intracranial recordings and stimulate with ultrasound at the same time. The small skull size of mice can also easily introduce standing waves at higher PRFs of ultrasound stimulation, which should be avoided to prevent artifacts and off target stimulation. Previous techniques for successful rat head-fixation have fully covered the skull with adhesives and the head-cap, maximizing surface area to keep a strong attachment (Roh et al., 2016; Schwarz et al., 2010). This is inadequate for tFUS however, as there must be space left on the top of the skull for a transducer to deliver the ultrasound energy. This means that a significant portion of the skull cannot be covered with the headcap or adhesives that do not easily allow much ultrasound energy through. In addition, alignment must be achieved between the chronically implanted electrode within the headcap and the delivered ultrasound energy.

In the present study, we develop an awake head-fixed rat model in which intracranial recordings can be taken simultaneously with aligned ultrasound stimulation. Using this model, we tested a range of PRFs and DCs to determine their effectiveness on RSU and FSU in the rat somatosensory cortex without the confound of anesthesia. By comparing these results to anesthetized ones, we demonstrate the cell-type specificity of tFUS and PRF sensitivity in an awake model, showing that this effect is not due to secondary anesthesia interactions. Our results provide a better understanding of the parameter space in awake rats, point towards the preferred parameters for inducing time-locked excitation, and help us better understand the mechanism of tFUS induced neural activation. This indicates the success of this model for performing intracranial recordings of tFUS neuromodulation without anesthesia, as well as its potential to help future tFUS awake experimentation.

## Methods

### Headpiece design

In order to record electrophysiological data from head-fixed rats while stimulating them with ultrasound, we developed a headpiece that would be compatible with our lab’s ultrasound transducer (Supplementary Figures 1A, 1B). Starting from the model developed by the Dudman Lab in 2014 (Osborne & Dudman, 2014) and successfully applied in several approaches (Belsey et al., 2020) we increased the size and added strips along the bottom to maximize surface area of headpiece connecting to the skull. A semicircle was cut out of the front to then leave room for the transducer to interface with the headpiece. Headpieces were 3-D printed from Formlabs “Clear Resin”.

Next, we designed arm bars that could be used to clamp the headpiece in place (Supplementary Figure 1C). The head of the arm bar was designed to align with the headpiece sides, while the bar portion was designed to fit into the ear bar slots in a stereotaxic setup. The arm bars were machined from 6061 aluminum to be both lightweight and sturdy.

### Awake Head-fixed Model Setup and Training

Ten adult male Wistar rats were used in this study, weighing approximately 250g at the beginning of our protocol. The animal study was conducted according to a protocol approved by the IACUC of Carnegie Mellon University. Rats were acclimated to human handling two weeks prior to surgery, and introduced during this time frame to their reward for future training, Froot Loops™.

On the day of surgery, the rats were anesthetized and given anti-inflammatory and anti-biotic medication. A square window was opened onto the skull, and a 2 mm burr hole was made in the S1 region of the rat brain (ML: -3 mm, AP: -2 mm, depth: 1 mm), where the electrode would be inserted, and a small hole in the dura was opened before the hole was filled with saline. Then, 10-12 1.19 mm Self-Tapping Bone Screws (19010-00, Fine Science Tools (USA) Inc., Foster City CA, USA) were inserted in approximately a rectangle behind the main burr hole, such that they would fit within the perimeter of the rectangular center of the headpiece as shown in Figure 1D. Screws were given two full twists for stability. The perimeter of the exposed skull was then covered with dental cement and the headpiece was glued down around the screws.

**Figure 1:**
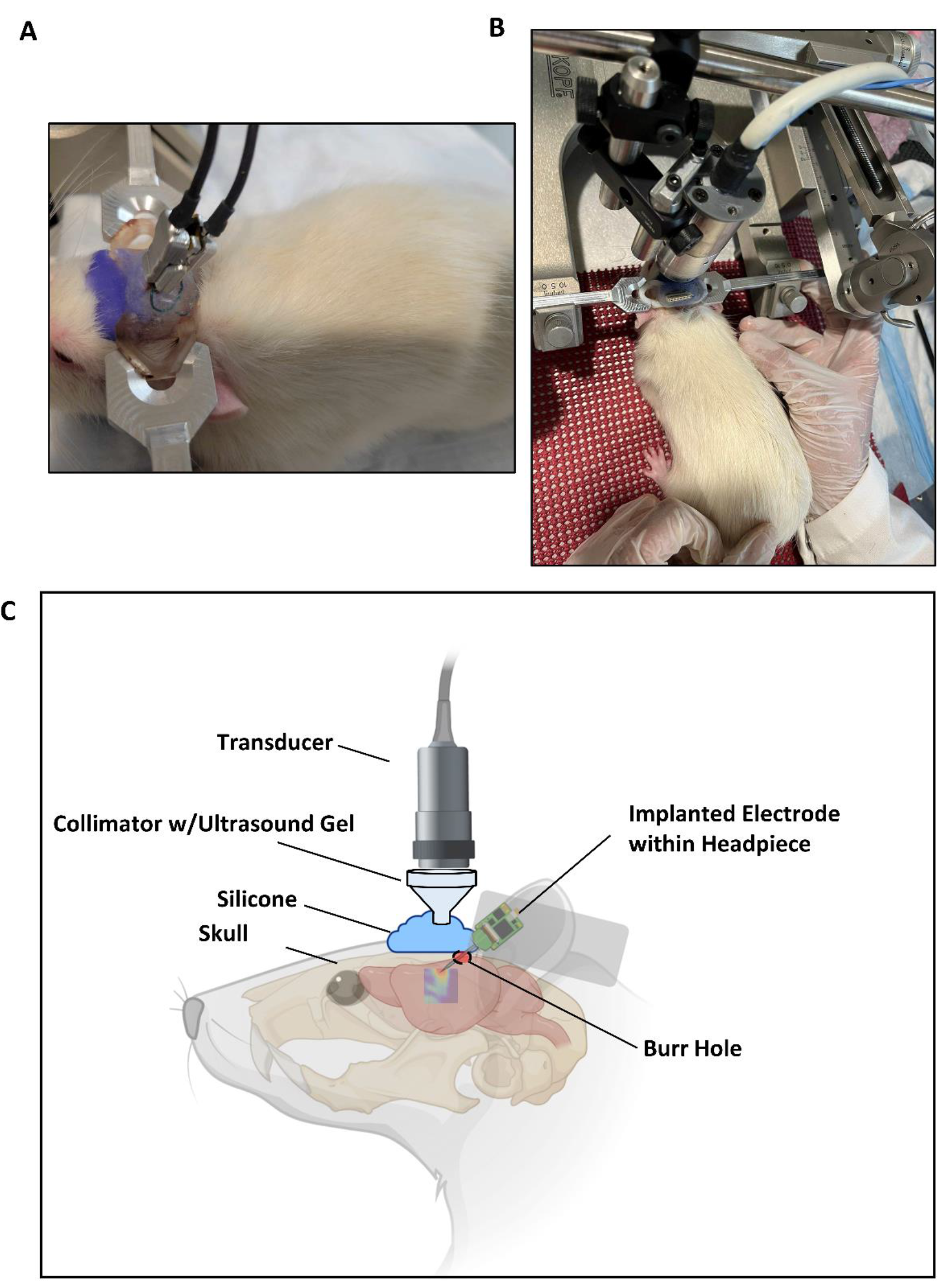
Awake Head Fixation Set-up. (A) A picture showing the headpiece on a rat in head- fixation, with the Zif-clip electrode and connector attached. (B) A picture showing the rat in head- fixation with the ultrasound transducer in position above. (C) A diagram showing the conformation of the transducer and ultrasound delivery relative to the rat.

At this point a 32-channel electrode was inserted, and grounding and reference wires attached to the skull screws (Supplementary Figure 1D, Figure 1A). The electrodes used were composed of 3 columns of channels on a 10 mm single shank, each recording site spaced out 50 microns from the next above and to the side (A1x32-Poly3-10mm-25s-177-Z32, Neuronexus, Ann Arbor, MI, USA). Zif-clip compatible electrodes were chosen for easy connection to a recording cable for awake recordings. The electrode was inserted at a 40 degree incidence angle such that the tip was closer to the front of the head. Body Double™ casting silicone (Smooth-On, Inc., Macungie, PA, USA) was placed over the inserted electrode to protect it and allow ultrasound transduction (Figures 1A and 1C). Finally, dental cement was used to secure the electrode in place and cover any remaining exposed area. Animals were allowed to recover for two weeks.

After recovery, training began by manually head-fixing the rats, holding them still within the experimental setup. Manual head-fixation was used initially so that the rats could struggle without any risk of damage to the head or headpiece. Rats were head-fixed for incrementally increasing times from 5 minutes up to 30 minutes each day across a week, five times a day with a break given in between. During each of these training sessions, Froot Loops™ were given during the rest periods. Training sessions were performed in the afternoon when the rats are least active, and human interaction was continued to keep the rats comfortable with their experimenter. After a week, rats could be switched to head-fixation using the arm-bars as shown in Figure 1A and 1B However, we observed that manual head fixation typically kept the rats calmer, so more startle-prone rats were recorded using manual-fixation. If a rat struggled significantly during training or recordings, a cloth wrap was used to help keep them still and calm during head-fixation. We observed that good surgical technique was very important to the success of the training, as any residual pain at the surgical site or rough edges from dental cement would result in a more stressed rat. Recordings were performed in a dimly lit recording booth with the door closed to minimize any sounds, which significantly calmed the rats.

### Ultrasound Setup

Ultrasound was delivered from our 128-element random array ultrasound transducer H276 from (*f*0: 1.5 MHz, -3dB focal size: 1.36 mm axially, 0.46 mm laterally), customized and manufactured by Sonic Concepts, Inc. (Bothell, WA, USA) (Supplementary Figure 2A). The transducer has a radius of acoustic aperture of 8.5 mm, and a 15 mm diameter of acoustic exit plane. Elements were arranged in a random distribution with 1 mm pitch (Supplementary Figure 2B). The transducer was driven with a Vantage 128 research ultrasound platform (Verasonics, Kirkland, WA, USA) using a DL-260 connector. An external power supply was used to ensure voltage stability during delivery of ultrasound bursts (QPX600DP, AIM-TTi, Cambridgeshire, UK). A collimator was designed to attach to the transducer to help with accurate targeting, and was 3D printed with VeroClear material. The collimator was filled with ultrasound gel (Aquasonic Ultrasound gel, parker Laboratories, Inc., Fairfield, NG, USA) and attached to the transducer before experiments. The collimator and transducer were pointed towards the targeted brain region where more ultrasound gel was placed to direct ultrasound energy smoothly through the collimator and skull, and into the brain. The Verasonics system was used to steer the ultrasound beam, with the collimator tip already aligned with the targeted brain region and depth targeting used to focus the beam through the collimator and into the brain at S1 (coordinates X=0, Y=0, Z=35).

Ex-vivo scanning was performed using a customized needle hydrophone-based 3D ultrasound pressure mapping system (HNR500, Onda Corporation, Sunnyvale, CA, USA), which allowed us to measure the necessary input voltage in order to account for any increased attenuation of ultrasound energy in the silicone layer over the skull. A rat skull sample was extracted from a euthanized rat subject complete with the silicone layer, and was placed between the H276 probe and the needle hydrophone in degassed water to mimic our experimental setup. We determined that an input voltage of 15 V resulted in a measured peak-to-peak ultrasound pressure of 114 kPa in the targeted region after penetrating the skull and silicone layer while using parameters of 1500 Hz PRF, a 67ms pulse duration (PD), and a 200 µs tone-burst duration (TBD) (Figure 2A and 2B).

**Figure 2:**
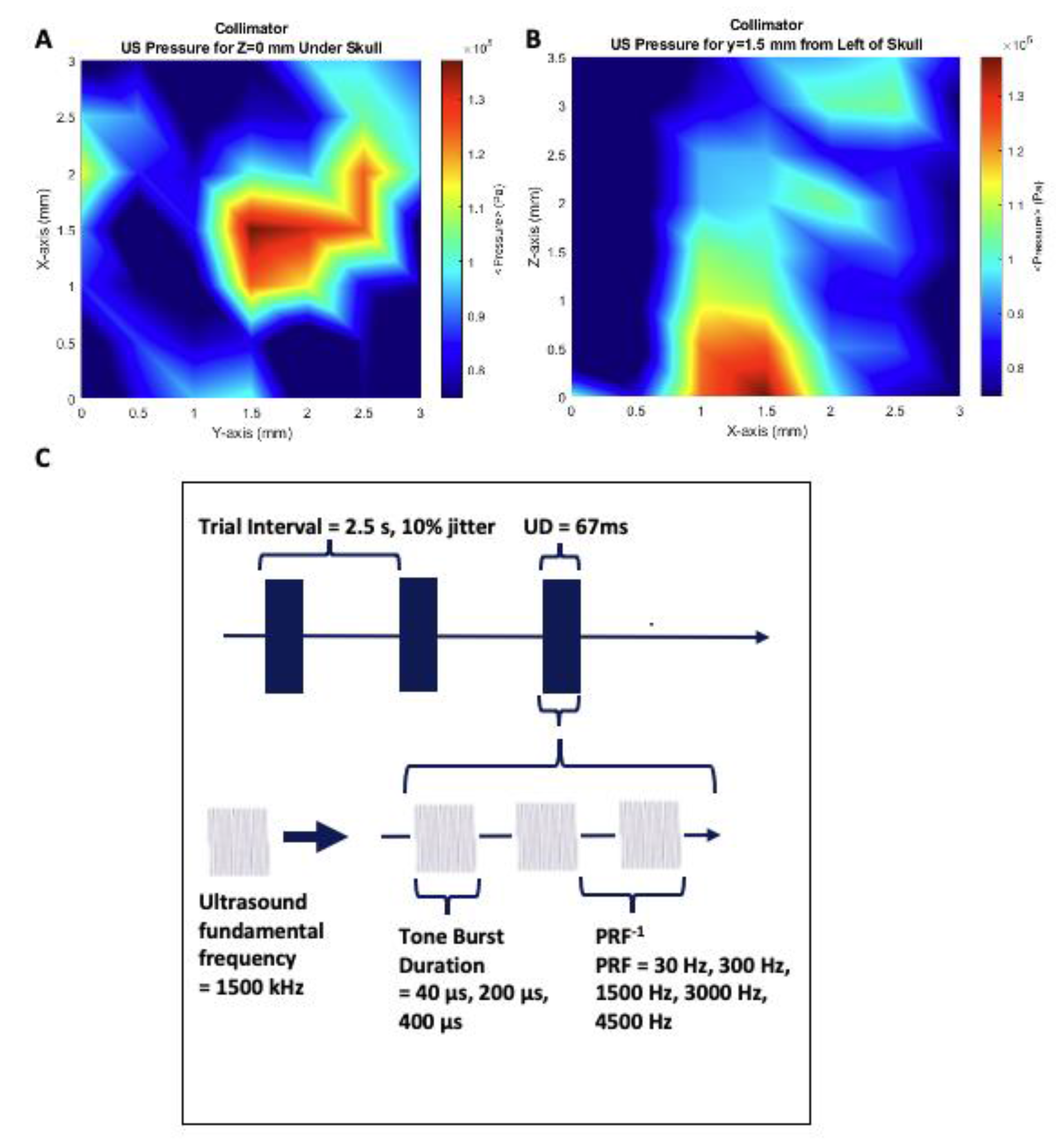
Ultrasound Field Characterization and Delivery Parameters. (A) Ex-vivo hydrophone peak-to-peak pressure amplitude z-axis scan at the expected depth of the targeted brain region. (B) Ex-vivo hydrophone peak-to-peak pressure amplitude y-axis scan at the expected position of the targeted brain region. (C) Diagram of the ultrasound delivery waveform parameters used in stimulation trials. The fundamental frequency of 1500 kHz on the bottom left is pulsed, at a variety of frequencies called the “Pulse Repetition Frequency” (PRF) and with varying Tone Burst Durations (TBD). A train of these pulses is delivered for 67ms, the ultrasound duration, in each trial. Trials are initiated every 2.5 seconds.

### Experimental Design

We repeated the 5 levels of PRFs previously tested by our group, 30 Hz, 300 H, 1500 Hz, 3000 Hz, and 4500 Hz, while using a TBD of 200 µs (Ramachandran et al., 2022; Yu et al., 2021) (Figure 2C). After we tested these parameters in our awake model, we repeated them in the same 10 rats under anesthesia. After PRF testing, we held PRF constant at 1500 Hz and tested tone burst durations of 40 µs, 200 µs, and 400 µs, resulting in duty cycles of 6%, 30% and 60%. Each parameter combination was tested by delivering 505 trials of ultrasound stimulation in a session, with one trial every 2.5 seconds. The ultrasound duration of each trial was 67 ms, and all trials used an *f0* of 1.5 MHz. 10% jitter was used in the inter-stimulus interval timing to minimize any potential brain adaptation to tFUS stimulation. Recordings for PRF tests were completed first, while rats were approximately 7 months old, 1 month after surgery. Anesthetized recordings were completed when rats were approximately 9 months old, and recordings for DC tests were completed at approximately 10 months. Recordings for a single rat for each set of parameters were completed on the same day, with either 2 or 3 recordings completed during each head-fixation session.

### Electrophysiology

Recordings were taken using the TDT Neurophysiology Workstation (24 kHz sampling frequency, Tucker-David Technologies, Alachua, FL, USA), and the neural trace was bandpass filtered between 300 Hz and 6 kHz for spike analysis. The recording system and ultrasound were grounded together to reduce noise. Characteristic raw neural trace responses to tFUS are shown in Figure 3A. Spike sorting was performed using the PCA-based spike classification software Offline Sorter (Plexon, Dallas, TX, USA). Waveform detection was performed using -3.5 standard deviations from the Mean of Peak Heights Histogram, and clustering was performed using a standard E-M (expectation-maximization algorithm) scan searching for up to three clusters in each channel, using prebuilt functions within the software. Due to the amount of noise in awake recordings due to movement artifacts, significant manual removal of artifacts was performed, as well as removal of cross-channel artifacts and high-amplitude artifacts. Detected neurons were rejected if their inter-spike interval (ISI) histograms showed a lack of neural refractory period (indicative of MUA), or periodic shape (indicative of electrical noise contamination).

**Figure 3:**
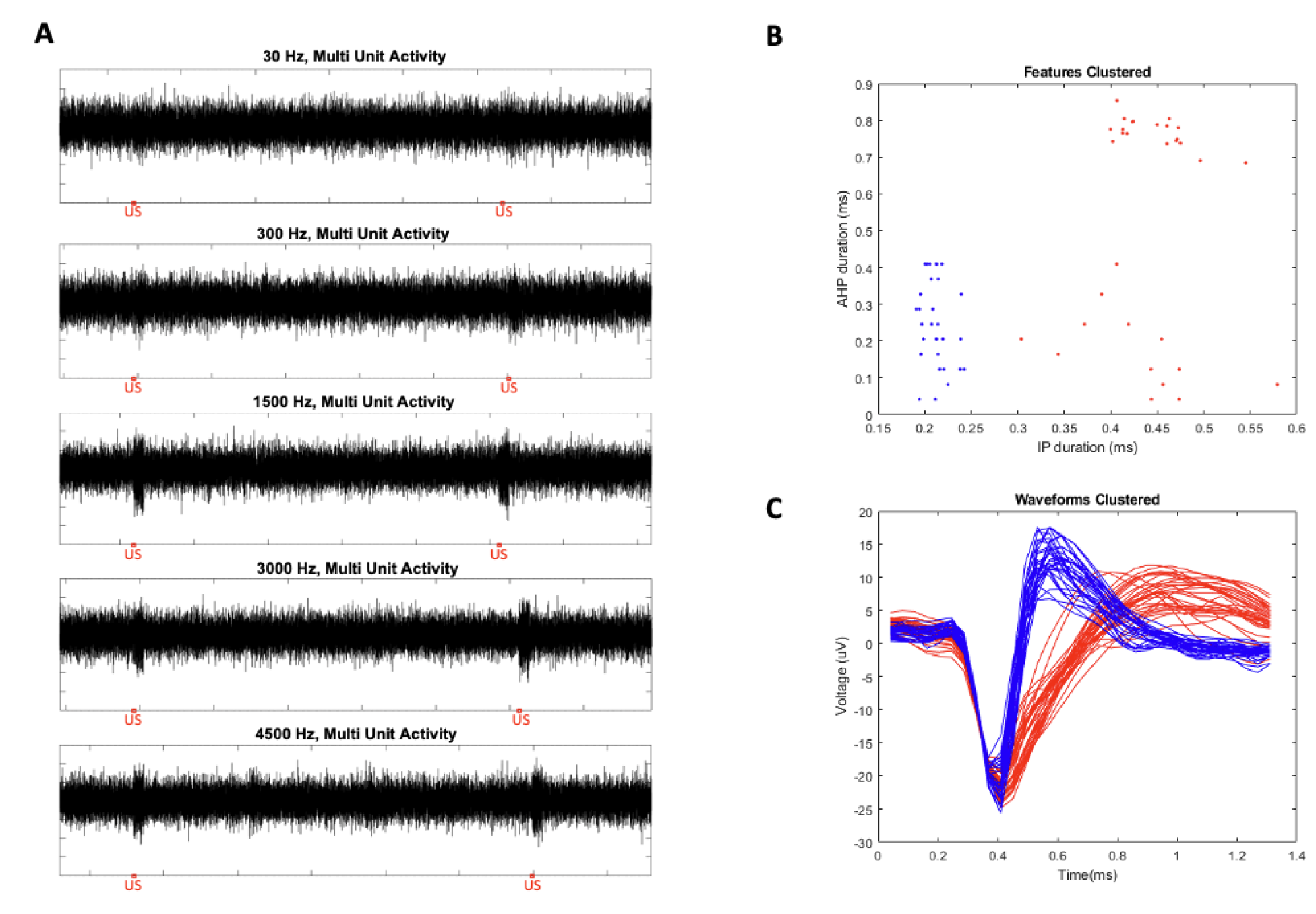
Multi-Unit Activity Response to tFUS and Cell-Type Sorting. (A) Examples of recorded multi-unit activity (MUA) from our awake head-fixed setup with tFUS delivered at five different PRFs (30 Hz, 300 Hz, 1500 Hz, 3000 Hz, 4500 Hz) in trials. (B) K-means clustering of detected neurons from one animals recordings, showing 28 FSU and 32 RSU, for 60 total. On the x-axis is plotted the initial phase (IP) of the action potential’s duration, and on the y-axis is the after- hyperpolarization (AHP) phase of the action potential’s duration. Blue dots were deemed to be FSU, and red dots were considered to be RSU. (C) Plotted average action potential waveforms for each of the clustered neurons shown in (B). Blue waveforms are FSU, while red waveforms are RSU.

After sorting, peri-stimulus time histograms (PSTHs) and raster plots were computed for each neuron in MATLAB R2019a (The MathWorks, Inc., Natick, MA, USA) using a combination of the FieldTrip toolbox (Oostenveld et al., 2011) and custom code. PSTHs were computed from 0.5 seconds prior to ultrasound stimulation, until 1.5 seconds afterwards, and were then divided by the number of trials (i.e., 500 after 5 are cut from the beginning of the recording), resulting in a histogram of average spikes per time bin for each neuron. Baseline average firing rate was computed from the 0.5 seconds before tFUS onset, and the response to tFUS was characterized as the normalized spike rate as a percent of baseline. Any PSTHs that were observed to have visibly fluctuated baseline periods, typically accompanied by periodic activity throughout the PSTH, were excluded from analysis and considered to be contaminated by artifacts.

We first characterized the previously studied time-locked response, defined as the response during the 67 ms ultrasound stimulation. We then studied delayed responses, defined as any response to tFUS after the 67 ms ultrasound duration.

### Cell Subtype Sorting

All neurons were sorted into RSU and FSU using custom code written in MATLAB based on the features of their action potential waveform. Specifically, the duration of the initial phase of the action potential, from onset to the first recrossing of baseline, and the duration of the after-hyperpolarization phase, from the first crossing of baseline to the recrossing of baseline were calculated for each neuron’s average waveform across each spike in a recording. K-means clustering was then used to sort the neurons based on these features (Figure 3B, 3C). In cases where neuronal features were not clearly in one cluster, spiking rate was used to further aid in sorting, and generally these were found to be RSU.

### Statistical Analysis

Responses of the different neuronal sub-groups within each condition in awake and anesthetized settings were compared with each other to test whether PRF and DC are significant factors, and to find which parameters elicit highest neuronal firing responses. Neuronal response data were tested for normality using the Shapiro-Wilt test, and were found to not have a normal distribution. Given this, the Kruskal-Wallis H test was used for group analysis of significance, and characterization was performed with *post hoc* Wilcoxon tests. The Bonferroni correction for multiple comparisons was used when examining p-values for significance. Parameters were recorded in a random order in each animal to mitigate any effect over time of recording.

## Results

### Differences in Response Latencies between Awake and Anesthetized Models

After confirming alignment of tFUS stimulation, we first examined the differences in neural responses to tFUS between awake and anesthetized recordings. We observed time locked responses to tFUS from RSU in both awake and anesthetized recordings, with an observable increase in firing rate within the 67 ms of tFUS (Figure 4C and 4E). In FSU we observed delayed responses to tFUS after the 67 ms duration of stimulation in both awake and anesthetized recordings, but observed that the response onset latencies of this response varied between the models (Figures 4D, 4F, and 4G). In the anesthetized model, delayed responses had an average onset latency of 0.24 s, while in the awake model they had an average onset latency of 0.51s (Figure 4I). The awake model also showed a wider range of onset latencies, and appears to have two peaks, one earlier within the anesthetized model’s distribution, and one later (Figure 4H). FSUs in the awake model generally showed a much wider variety of responses than in the anesthetized model, with two examples of responses shown in Figures 4F and 4G. Figure 4F shows the higher latency response that was most commonly observed, while Figure 4G shows an earlier latency response more similar to that observed in the anesthetized model. Interestingly, Figure 4G also shows slight but significant inhibition time locked with tFUS. Comparing the distributions using the Wilcoxon rank-sum test, we found that delayed response onset times varied significantly between awake and anesthetized models (p < 0.001).

**Figure 4:**
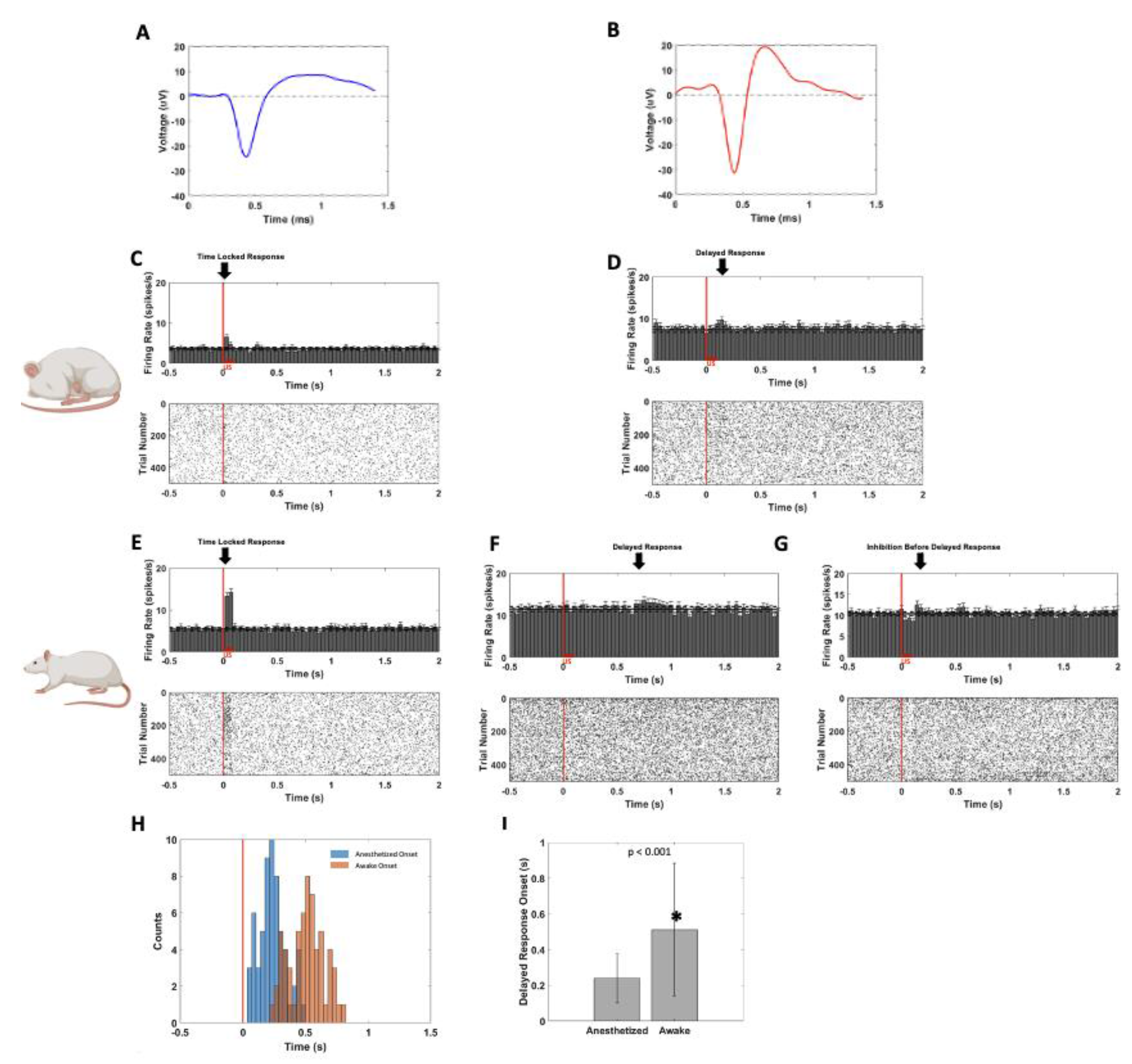
Varying Latencies in Cell-type Specific Neuronal Spiking Responses to tFUS in Awake and Anesthetized Models. (A) A characteristic waveform of a regular spiking unit (RSU) from acquired recordings. (B) A characteristic waveform of a fast spiking unit from acquired recordings. (C), (D), (E), (F), (G) Representative peri-stimulus time histogramas (PSTHs, bin size: 35.7 ms, n=505 trials for each time bin) and raster plots of responses to trials of 1500 Hz PRF 30% DC tFUS from (C) an RSU in the anesthetized model, (D) an FSU in the anesthetized model, (E) an RSU in the awake model, and (F) and (G) two different FSU in the awake model. The vertical red line shows the tFUS onset. (H) Plotted histograms of delayed response onset latencies in the anesthetized model (blue) and the awake model (orange). The horizontal red line shows the tFUS onset. (I) A bar plot showing the difference in average delayed response onset latency between the awake and anesthetized model. Significance was characterized with a t-test, and the groups were found to be significantly different with p <0.001.

### Awake vs Anesthetized Model Sensitivity to PRF

We tested PRFs of 30 Hz, 300 Hz, 1500 Hz, 3000 Hz, and 4500 Hz while holding the TBD at 200 µs in both the awake and anesthetized model, first examining the time-locked response, and then the delayed response in order to determine whether cell type selective responses and PRF sensitivity are maintained in an awake model. Examining the time-locked response from RSU in the awake model using Kruskal-Wallis ANOVA, we find that PRF is a significant factor (p < 0.001). Post hoc tests show that 30 Hz and 300 Hz PRF induced responses lower than all higher PRFs (p< 0.001). 1500 Hz, 3000 Hz, and 4500 Hz showed no significant differences (Figure 5A). The anesthetized model shows similar results, with ANOVA showing that PRF is significant (p<0.01), 30 Hz and 300 Hz being significantly lower than the others (p <0.001), and no significant difference between the highest three PRFs (Figure 5C). Two-way ANOVA additionally finds that anesthesia is a significant factor in the time-locked response from RSU (p <0.01). FSU in both the awake and the anesthetized model show no significant differences due to PRF levels (Figures 5B and 5D).

**Figure 5:**
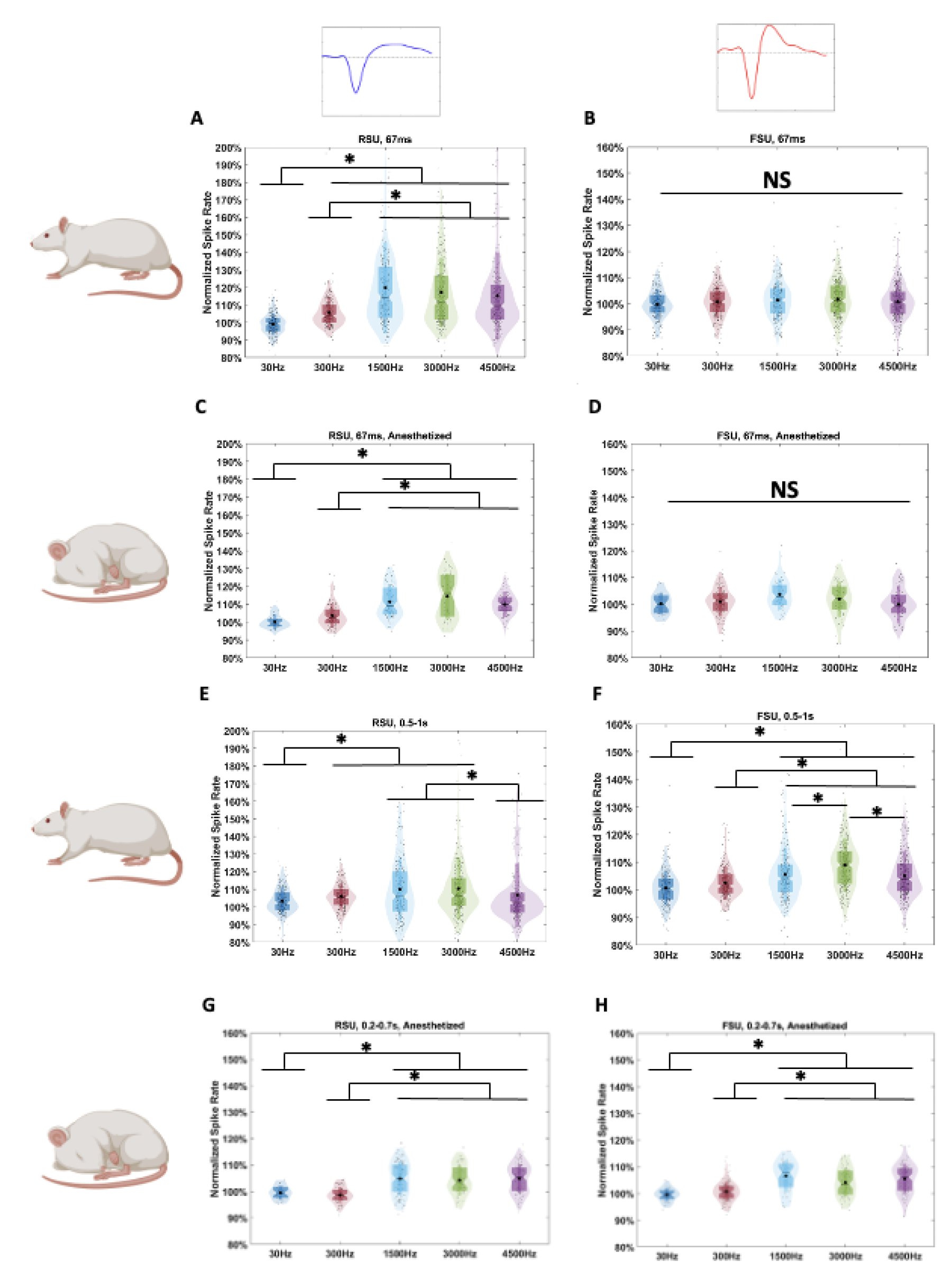
Group Analysis of Cell-type Specific Responses to tFUS with Varying PRF in Awake and Anesthetized Models. (A), (B), (C), (D), (E), (F), (G), and (H) all show violin plots of the average normalized spike rate induced by tFUS within each parameter over 505 trials in the specified time range compared to baseline, calculated from the 0.5 seconds before stimulation. Each small black dot represents a neuron from a recording under the specified parameters. The black asterisks show the mean of each group. The darker color figures are traditional box plots with the ends of the boxes marking the 1^st^ and 3^rd^ quartiles and the middle line marking the median. Notches mark the 95% confidence intervals on the median. The narrow rectangle edges mark one standard deviation to each side of the mean of the group. The lighter colored figures are the probability distribution of the data, with the ends of the figure marking the 1^st^ and 99^th^ percentile. Statistics are performed using the Kruskal-Wallis two-sided one-way ANOVA on ranks, with post hoc Wilcoxon tests with the Bonferroni correction for multiple comparisons used afterwards. (A) Time locked responses from RSU in the awake condition. Significantly different responses to different PRFs are observed. (B) Time locked responses from FSU in the awake condition. No significant differences are observed. (C) Time locked responses from RSU in the anesthetized condition. Significantly different responses to different PRFs are observed. (D) Time locked responses from FSU in the anesthetized conditions. No significant differences are observed. (E) Delayed responses (0.5-1 s) from RSU in the awake condition. Significantly different responses to different PRFs are observed. (F) Delayed responses (0.5-1 s) from FSU in the awake condition. Significantly different responses to different PRFs are observed. (G) Delayed responses (0.2-0.7 s) from RSU in the anesthetized condition. Significantly different responses to different PRFs are observed. (H) Delayed responses (0.2-0.7 s) from FSU in the anesthetized conditions. Significantly different responses to different PRFs are observed.

Next, we considered the delayed responses. Using ANOVA, PRF was found to be a significant factor for both RSU and FSU in both awake and anesthetized settings (Figures 5E, 5F, 5G, and 5H). Results from RSU in the awake setting show that 1500 Hz and 3000 Hz induce the highest responses (p < 0.001), showing a decrease at higher frequencies in the delayed response from RSU (Figure 5E). In the anesthetized setting, the results are more similar to the time-locked response, with 1500 Hz, 3000 Hz, and 4500 Hz all resulting in the highest responses compared to 30 Hz and 300 Hz (p < 0.001, Figure 5G). FSU in the awake setting show the strongest responses to 3000 Hz PRF, with significant differences among all PRFs (p<0.001, Figure 5F). In the anesthetized model, FSUs still show fewer significant differences, with similar results as in the RSU’s anesthetized delayed response, showing the highest responses to 1500 Hz, 3000 Hz, and 4500 Hz compared to 30 Hz and 300 Hz (p < 0.001, Figure 5H). Using two-way ANOVA, we find that anesthesia is a significant factor in the delayed response for both RSU (p < 0.001) and FSU (p < 0.01). We observe overall that in the awake model more high responding outliers are present, especially in RSU, and tails of the data distributions overall extend higher. We also observe that in general 1500 Hz, 3000 Hz, and 4500 Hz produce the significantly highest responses when PRF sensitivity is observed, with no significant differences between them. Although it is not statistically significant, we find that 1500 Hz produces the highest mean time-locked response (Figure 5A).

### Awake Model Sensitivity to Duty Cycle

Following the observation that 1500 Hz was consistently capable of producing excitation, and that it induced the highest mean response (although not significantly different from other high PRFs in most cases), we tested the effect of varying the DC by changing TBD while holding PRF constant at 1500 Hz on neural responses. In the time-locked responses, RSU showed a significantly lower response to 6% DC than 30% or 60% (p < 0.01, Figure 6A). In the delayed response however, RSU responded with significantly higher spiking rates in 30% DC than those in both 6% and 60% (p < 0.001, Figure 6C), with significant differences observed between 6% and 60% as well (p < 0.01). FSU showed no significant effect from varying DC in the time-locked response (Figure 6B), but in the delayed response 60% was found to induce a significantly higher response than 6% (p < 0.001, Figure 6D). Based on mean values, 60% appears to produce the strongest time-locked response, although it produces the lowest response from the delayed response.

**Figure 6:**
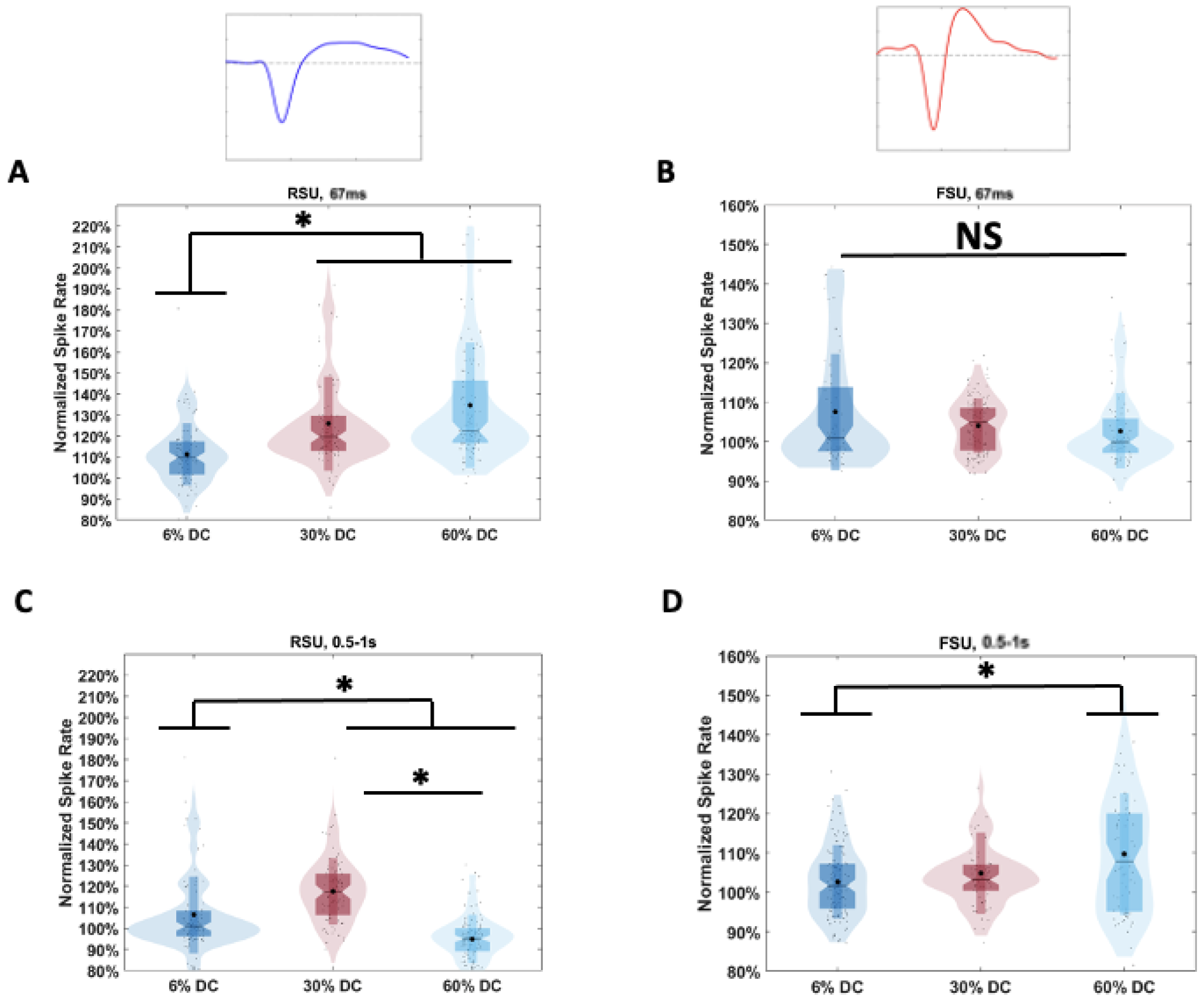
Cell-type Specific Responses to Varying Duty Cycle of tFUS in the Awake Model. (A), (B), (C), and (D) show violin plots for each condition of the average normalized spike rate induced by tFUS over 505 trials in the specified time range compared to baseline, calculated from the 0.5 seconds before stimulation. Statistics are performed using the Kruskal-Wallis two-sided one-way ANOVA on ranks, with post hoc Wilcoxon tests with the Bonferroni correction for multiple comparisons used afterwards. (A) Time locked responses to DC among RSU. 6% DC results in a significantly lower response than 30% or 60%. (B) Time locked responses to DC among FSU. No significant differences are observed (C) Delayed responses (0.5-1 s) to DC among RSU. Significantly different responses are observed between all DCs. (D) Delayed responses (0.5-1 s) to DC among FSU. 6% and 60% DC show significantly different levels.

## Discussion

In this study we have developed an awake head-fixed rat model to simultaneously deliver ultrasound stimulation and take electrophysiological recordings. Using this novel animal model, we have investigated a variety of ultrasound parameters to test our hypothesis that tFUS would induce cell-type specific neuronal responses to both PRF and DC variation. Our study of the ultrasound parameter space without the confound of anesthesia provides a better understanding of the mechanisms behind tFUS neuromodulation in an awake animal model. The present results show how PRF and DC levels affect tFUS responses, and give us more evidence pointing towards the mechanisms of tFUS stimulation in different cell-types.

### Awake Head-fixed Model

Our development of an awake head-fixed rat model was successful in allowing us to take electrophysiological recordings with high spatial and temporal specificity in an awake rat while delivering ultrasound stimulation. Our results show that this model successfully achieved alignment between the ultrasound beam and the recorded brain region, allowing us to record spiking activity in an awake rat model responding to ultrasound for the first time to the best of our knowledge. This setup was simple to implement as it did not require any specialized transducer and used a regular stereotaxic setup. The headpieces themselves were 3D printed at low-cost. Weaknesses of this model as currently developed are mainly noise management and ensuring that targeting is done the same way for each recording. To ensure minimal noise, the recording room must be electrically isolated, and grounding during surgery for each animal needs to be performed carefully. Targeting relies on carefully implanting the electrode in the same position relative to the headpiece each time, and aligning the transducer to the headpiece the same way each time. This is generally the protocol used in acute recordings as well, but it means that there are likely at least slight differences in alignment between recordings, which have the potential to result in varying degrees of excitation from tFUS, and may be a source of the variation in responses that we see in our results. Future iteration on this model may benefit from attempting to improve the consistency of targeting. Despite this, we consider this model a beneficial new tool for future testing of neural responses to array-based tFUS neuromodulation in an awake setting for rat models, making future behavioral testing simpler as well.

### Awake Model Cell Type Specificity and Parameter Sensitivity

We recorded awake neural responses to tFUS in the S1 cortex, testing a variety of different PRF levels and DCs. The results provide *in* vivo evidence that RSU respond in a time-locked manner that is sensitive to increases in PRF in awake animals. We also observe this under anesthesia, aligning with our previous studies (Yu et al., 2021). FSU, unlike RSU, show no significant time-locked response to PRF changes in either the awake or anesthetized recordings. This demonstrates that the cell-type differentiated effect is not due to an anesthesia confound. We then examined delayed responses to tFUS, observed in both RSU and FSU, although smaller than time-locked responses. Both RSU and FSU delayed responses are sensitive to PRF level, but instead of strictly increasing with PRF show a decline at higher levels such as 4500 Hz. That we see PRF sensitivity in the delayed response shows that the same tFUS mechanism of excitation may be involved in both time-locked and delayed responses in both RSU and FSU.

To determine effects of DC variation, we then varied the TBD, effectively altering the DC while holding PRF constant at 1500 Hz. RSU’s time locked response responded stronger to higher PRFs, while its delayed response was strongest at 30% DC in the middle range. By contrast, FSU’s delayed response was strongest at 60% DC, once again showing no significant time-locked response. This demonstrates that changing DC may also impact cell-type specificity in induced excitation. The differences in preferred parameters between time-locked and delayed responses may indicate that the way the tFUS mechanism works in the different cell types and between the time-locked and delayed responses varies.

Sensitivity to both PRF and DC suggests that the dynamic acoustic radiation force (ARF) may be a significant mechanism of ultrasound neuromodulation at play. ARF as it applies to tFUS is the force resulting from attenuation of momentum of the ultrasound wave into the brain tissue. Mechanical force such as this stretches cell membranes and modulates ion channel conformations as well as their fluid environments, altering membrane conductance and thus spiking probabilities. The magnitude of the ARF is calculated using the spatial-peak temporal-average intensity (ISPTA), which depends on PRF and DC/TBD (Yu et al., 2021).

As discussed in the introduction, different neuronal subtypes express different ion channel distributions as well as different overall shapes and dendritic arbor alignments within the cortex. Our group previously hypothesized that these differences were the cause of the cell-type specific sensitivity to PRF changes, in relation to ARF effects. In this study, we confirm that this cell type specificity does not come from cell-type specific anesthesia effects, providing further evidence that ARF is the mechanism for this differentiation.

### Differences between Awake and Anesthetized models

We found that anesthesia was not a confound to the cell-type specific effect, but did observe differences between responses in the awake and anesthetized setting. RSU responded similarly in both model although more strongly in the awake setting, but FSU exhibited more varied behavior. Although previous analysis has focused on time-locked responses to tFUS, delayed responses have been observed before. PSTHs formed from MUA signal in Tufail *et al*’s 2010 paper studying anesthetized mice shows a smaller peak at approximately 250 ms post stimulation (Tufail et al., 2010). Calcium imaging in Murphy *et* al’s 2022 paper investigating cell-type specific responses to tFUS found that PV+ inhibitory interneurons (likely FSU) exhibited a slow increase in activity 1-3 seconds after stimulation (Murphy et al., 2022b). We previously observed no delayed responses in FSU but while using a ketamine/xylazine cocktail anesthesia (Yu et al., 2021). This anesthesia method was found to significantly modulate responses to PRF level and cell type responses relative to later isoflurane recordings. It was also specifically observed that fewer FSU were recorded overall under ketamine/xylazine compared to isoflurane, indicating that it may have a particularly dampening effect on FSU. Our group’s more recent research has indeed observed delayed responses in FSU under isoflurane anesthesia (Gao et al, 2024). Even under isoflurane anesthesia however, we did not observe the later delayed responses observed in our awake model. Additionally, our awake model showed some FSU to have a time-locked inhibitory response in addition to a later excitatory delayed response.

The mechanism behind the delayed responses as opposed to the time-locked responses is not fully clear. Murphy *et al* hypothesized that PV+ cells from their calcium imaging have a different mechanism causing them to respond to tFUS in a more delayed manner, or that it may be a polysynaptic effect. Our own group has hypothesized that these delayed responses are most likely due to CTC (cortico-thalamo- cortical) circuit activation and feedback (Gao et al, 2024). It has been demonstrated that CTC connectivity is significantly reduced by anesthesia in rats (Gao et al., 2017; Liang et al., 2015). If circuit activity is the cause of the delayed responses, this may explain why we observe a difference in the awake model.

Another potential explanation is that the higher latency responses observed from FSU in the awake model may be due to a different mechanism of ultrasound activation which is blocked under anesthesia. This mechanism may be more in line with the slow PV+ activation observed by Murphy *et al* also in awake animals (Murphy et al., 2022b). Under our theory of an ARF mechanism, the ion channel distribution and geometrical shape of RSU may result in quick elicitation of activity, while the ion channel distribution and shape of FSU may result in a slower elicitation that can be blocked by anesthesia effects on those ion channels.

### Future Directions and Confounds

One limitation in our study is the relatively low number of animals used. Unlike other methods of stimulation, ultrasound is known to have particularly variable effects. Trial to trial, stimulation is unlikely to result in the same amount of excitement each time, and inter-subject differences appear to be relatively high, due to effects of different brain morphology or even skull shape. Beyond this, orientation of different neurons to the ultrasound wave potentially affects how they are stimulated, meaning that different neurons within a recording may be effectively experiencing different ultrasound delivery, even when they are targeted directly with the same pressure levels. Additionally, as previously mentioned, targeting intrinsically came with some variation although minimized as much as possible. Numbers in this study were limited due to the challenge of training and maintaining AHF model animals over months, but future studies may wish to use more animals to clarify differences between parameter levels.

Another limitation in our study was the range of parameters tested. In our tests of PRF, we hold the TBD constant, which means that DC in fact varied with PRF. After this, we held PRF constant and tested different TBDs, changing the DC. Our previous study determined that the cell-type specific effect was also present while holding DC constant and varying PRF and TBD, but this was not repeated here. We additionally kept the fundamental frequency, intensity level, and ultrasound duration constant, which future studies may wish to investigate further. Here we focused on PRF and TBD/DC so that we could compare to our previous anesthetized study, and because we are particularly interested in the time-locked excitation effects potential to induce plasticity within a certain delivery paradigm (Ramachandran et al., 2022).

A potential confound to our findings are that the PRF and DC data are taken a few months apart. In general, implanted electrodes are expected to see a decrease in signal quality over time. In our recordings we made sure to confirm that we could still see neural activity and ultrasound effects in our recordings before proceeding, but this decrease in quality may still have had some effect on our results when comparing recordings taken at 7 months and 10 months, as could have the increased age of the rats. Another confound comes from the vibratory nature of ultrasound. It is possible for ultrasound to induce indirect auditory activation (Guo et al., 2018; Sato et al., 2018). In our previous anesthetized study with similar setup (Yu et al., 2021) we tested several sham conditions of ultrasound delivery that would induce similar noise or skull vibration, and found that they did not elicit any significant response at any parameters. Based on this, we consider this not to be a confound to these results, but did not directly test for it in this study. Another common concern in intracranial recording tests of tFUS is that the vibration may make the electrode tip itself vibrate, causing local effects. A previous study however suggested that electrode tip movement is not significant under pressure levels of 131 kPA (M. G. Kim et al., 2021) . Since our stimulation is expected to elicit pressure levels of 114 kPA, we consider this not to be a concern in this study.

## Conclusions

This study confirms the cell-type specific and parameter dependent qualities of ultrasound in an awake rodent model. We observed that RSUs responded to tFUS in a time-locked manner, while FSUs responded at a delayed latency, that was modulated by anesthesia. Both these responses showed sensitivity to PRF and TBD/DC. These results provide further evidence that ARF is a key mechanism behind tFUS neuromodulation. Although our findings indicate that anesthesia was not a major confound to the observations of these characterizations of tFUS-neuron interactions, we also observe that anesthesia significantly modulates the response to tFUS, and that FSU delayed responses show modulated dynamics under anesthesia. Future investigation of tFUS effects on circuit dynamics or different cell types should consider this confound of anesthesia. Our developed AHF model for tFUS in a rat model will allow for intracranial electrophysiological investigation into tFUS without anesthesia in the future.

## Data availability

Experimental data supporting the findings will be made available in a public online repository when the paper is published.

## CRediT authorship contribution statement

**Sandhya Ramachandran**: Writing – review & editing, Writing – original draft, Formal analysis, Data curation, Investigation, Conceptualization. **Huan Gao**: Writing – review & editing, Data curation, Investigation. **Eric Yttri**: Writing – review & editing, Methodology. **Kai Yu**: Writing – review & editing, Methodology, Investigation, Supervision, Conceptualization. **Bin He**: Writing – review & editing, Investigation, Supervision, Project administration, Conceptualization.

## Acknowledgments

We thank Dr. Min Gon Kim for useful discussions on experimental procedures. Some images in Figures and Supplementary Figures were created with <u>BioRender.com</u>.

---

**Supplementary Figure 1.**
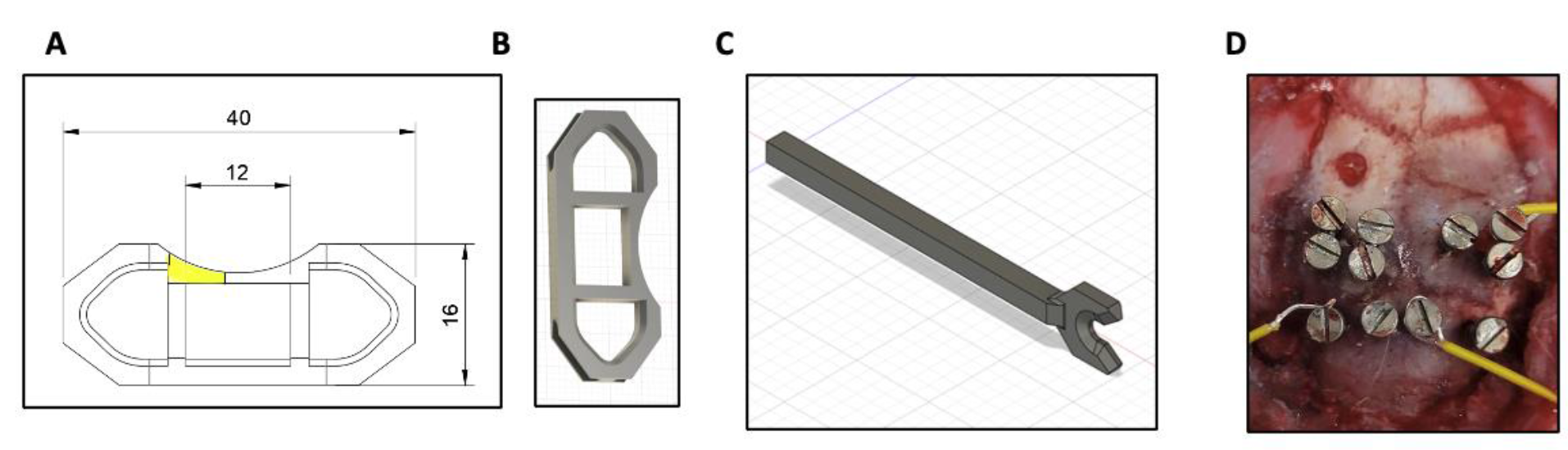
: Design of the Headfixation Headpiece. (A) A design blueprint for the headpiece, showing the dimensions of each segment in millimeters. (B) A 3D model of the headpiece. (C) A 3D model of the arm bar used for head-fixation. (D) A picture taken during the surgical protocol showing the relative positioning of the burr hole and skull screws, with grounding and reference wires shown attached to the screws.

**Supplementary Figure 2.**
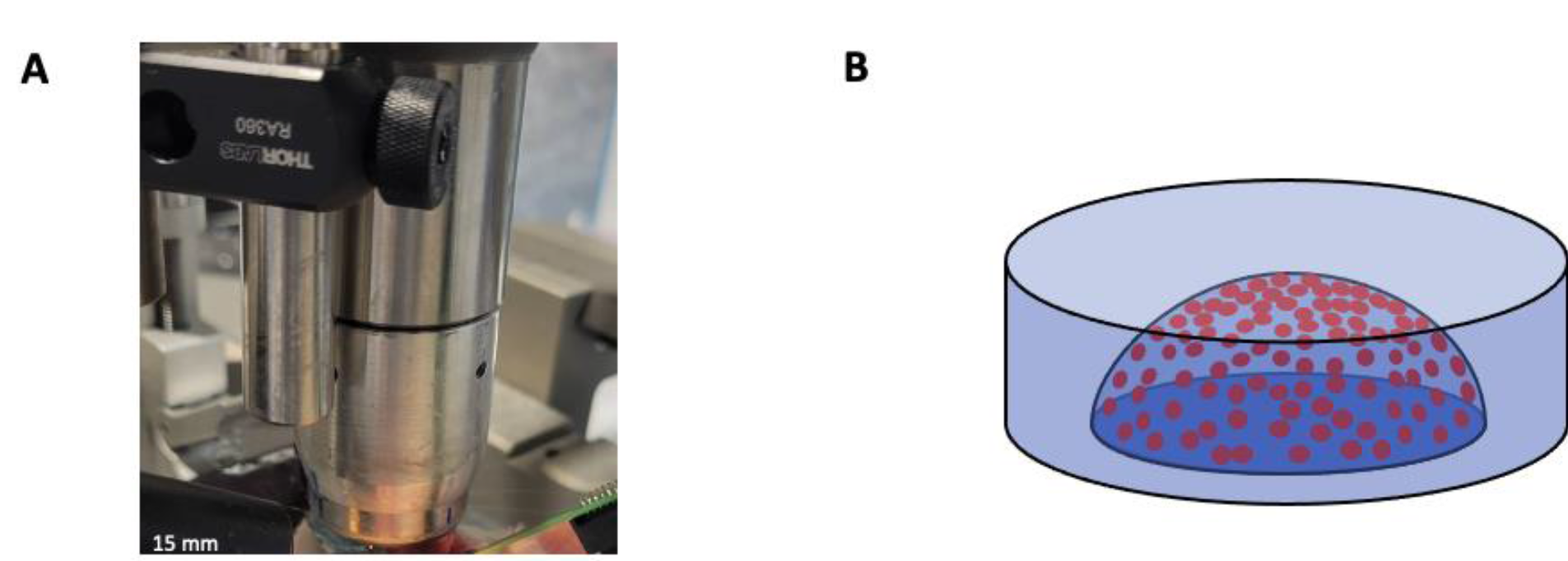
: Ultrasound Transducer Design. (A) A picture of the H276 128-element random array transducer. (B) Model of the H-276 element positioning.

